# Multiscale computer modeling of spreading depolarization in brain slices

**DOI:** 10.1101/2022.01.20.477118

**Authors:** Craig Kelley, Adam JH Newton, Sabina Hrabetova, Robert A. McDougal, William W Lytton

## Abstract

Spreading depolarization (SD) is a slow-moving wave of neuronal depolarization accompanied by a breakdown of ion concentration homeostasis, followed by long periods of neuronal silence (spreading depression), and associated with several neurological conditions. We developed multiscale (ions to tissue slice) computer models of SD in brain slices using the NEURON simulator: 36,000 neurons (2 voltage-gated ion channels; 3 leak channels; 3 ion exchangers/pumps) in the extracellular space (ECS) of a slice (1 mm sides, varying thickness) with ion (K^+^, Cl^−^, Na^+^) and O_2_ diffusion and equilibration with a surrounding bath. Glia and neurons cleared K^+^ from the ECS via Na^+^/K^+^ pumps. SD propagated through the slices at realistic speeds of 2–4 mm/min, which increased by as much as 50% in models incorporating the effects of hypoxia or propionate. In both cases, the speedup was mediated principally by ECS shrinkage. Our model allows us to make testable predictions, including: **1**. SD can be inhibited by enlarging ECS volume; **2**. SD velocity will be greater in areas with greater neuronal density, total neuronal volume, or larger/more dendrites; **3**. SD is all-or-none: initiating K^+^ bolus properties have little impact on SD speed; **4**. Slice thickness influences SD due to relative hypoxia in the slice core, exacerbated by SD in a pathological cycle; **5**. SD and high neuronal spike rates will be observed in the core of the slice. Cells in the periphery of the slice near an oxygenated bath will resist SD.

**Significance:** Spreading depolarization (SD) is a slow moving wave of electrical and ionic imbalances in brain tissue and is a hallmark of several neurological disorders. We developed a multiscale computer model of brain slices with realistic neuronal densities, ions, and oxygenation. Our model shows that SD is exacerbated by and causes hypoxia, resulting in strong SD dependence on slice thickness. Our model also predicts that the velocity of SD propagation is not dependent on its initiation, but instead on tissue properties, including the amount of extracellular space and the total area of neuronal membrane, suggesting faster SD following ischemic stroke or traumatic brain injury.

## Introduction

Spreading depolarization (SD), is a slow-moving (1.7–9.2 mm/min), long-lasting (minutes) wave of neuronal depolarization accompanied by a breakdown in homeostatic maintenance of intra- and extracellular ion concentrations, and associated with reduced neuronal activity (spreading depression) [Woitzik et al., 2013, Newton et al., 2018, Cozzolino et al., 2018, Dreier, 2011]. SD has been observed in a number of species, can be elicited experimentally both *in vivo* and in brain slices, and has been implicated in several neurological conditions, including ischemia, migraine, traumatic brain injury, and epilepsy [Cozzolino et al., 2018]. SD is difficult to detect in humans noninvasively [Drenckhahn et al., 2012, Hartings et al., 2014, Hofmeijer et al., 2018, Zandt et al., 2015], making it important to study SD in experimental preparations and computer simulation to better understand its role in human disease, and possible treatments.

SD has been studied in brain slices from a wide range of species and brain regions, including neocortex, hippocampus, brainstem, and retina [Martins-Ferreira et al., 2000, Balestrino et al., 1988, Aitken et al., 1998, Müller and Somjen, 1998, Andrew, 2016, Devin Brisson et al., 2013, Hrabe and Hrabetova, 2019]. It can be triggered experimentally by various means, including electrical stimulation; mechanical insult; K^+^ and ouabain application [Leao, 1944, Bures et al., 1974, Balestrino, 1995, Aitken et al., 1998, Joshi and Andrew, 2001]. SD can be facilitated by applying propionate to the slice [Tao et al., 2002, Hrabe and Hrabetova, 2019].

SD is intimately related with hypoxia in both slice and *in vivo*. Ischemia plays a complex role in ischemic diseases: hypoxia can initiate SD which then can contribute to the extent of the ischemic penumbra [Nedergaard, 1988] SD can itself trigger ischemia. The scintillating scotomata of classical migraine is believed to be an SD wave and increases risk of stroke[Øie et al., 2020] ; complicated migraine is caused by more severe ischemia, which produces more pronounced and long-lasting deficits [Santos et al., 2012]. In *in vivo* experiments, SD also led to hypoxia [Takano et al., 2007, Piilgaard and Lauritzen, 2009]. To begin examining this complexity in our simulations, we identified 3 types of hypoxia related to SD, comparable to different slice experiments. 1. We induced hypoxia to in turn induce depolarization, a phenomenon that has been called *hypoxic SD-like depolarization* (HSD) [Balestrino et al., 1988, Aitken et al., 1991]. 2. We used the “classical” SD-initiation protocol (adding a K^+^ bolus of to a slice) under hypoxic conditions, but before HSD had initiated, in order to compare to SD with an oxygenated bath. 3. We looked at how SD induced hypoxia as perfusion from the bath failed to keep up with the metabolic demand of overworked Na^+^/K^+^ pumps.

In this paper, we used multiscale computational modeling of SD to relate the microscopic levels of ion and O_2_ diffusion, channels, and pumps to the neuronal level of cell spiking up to the macroscopic level of tissue activation patterns (Fig. 1). Our baseline model was composed of 36,000 biophysically detailed point-neurons in an ECS of a square slice (1 mm sides, 400 *μ*m thick) with O_2_ perfusion and ion flux with a surrounding bath where relevant concentrations are held constant at their baseline values. We simulated SD in both perfused and hypoxic slices. Our model showed that SD speed was augmented by propionate and hypoxia and suggested that changing the ECS was the principle mechanism through which they influence SD. We predicted that SD speed changes with slice thickness due to core hypoxia and increases with the total neuronal surface area in the tissue. SD speeds in all conditions were enhanced by hypoxia. We also predicted that increasing the size of the ECS relative to the tissue will inhibit SD. Finally, we identified a depth-dependent relationship with greater SD propagation through core of the slice compared to the periphery.

**Figure 1:**
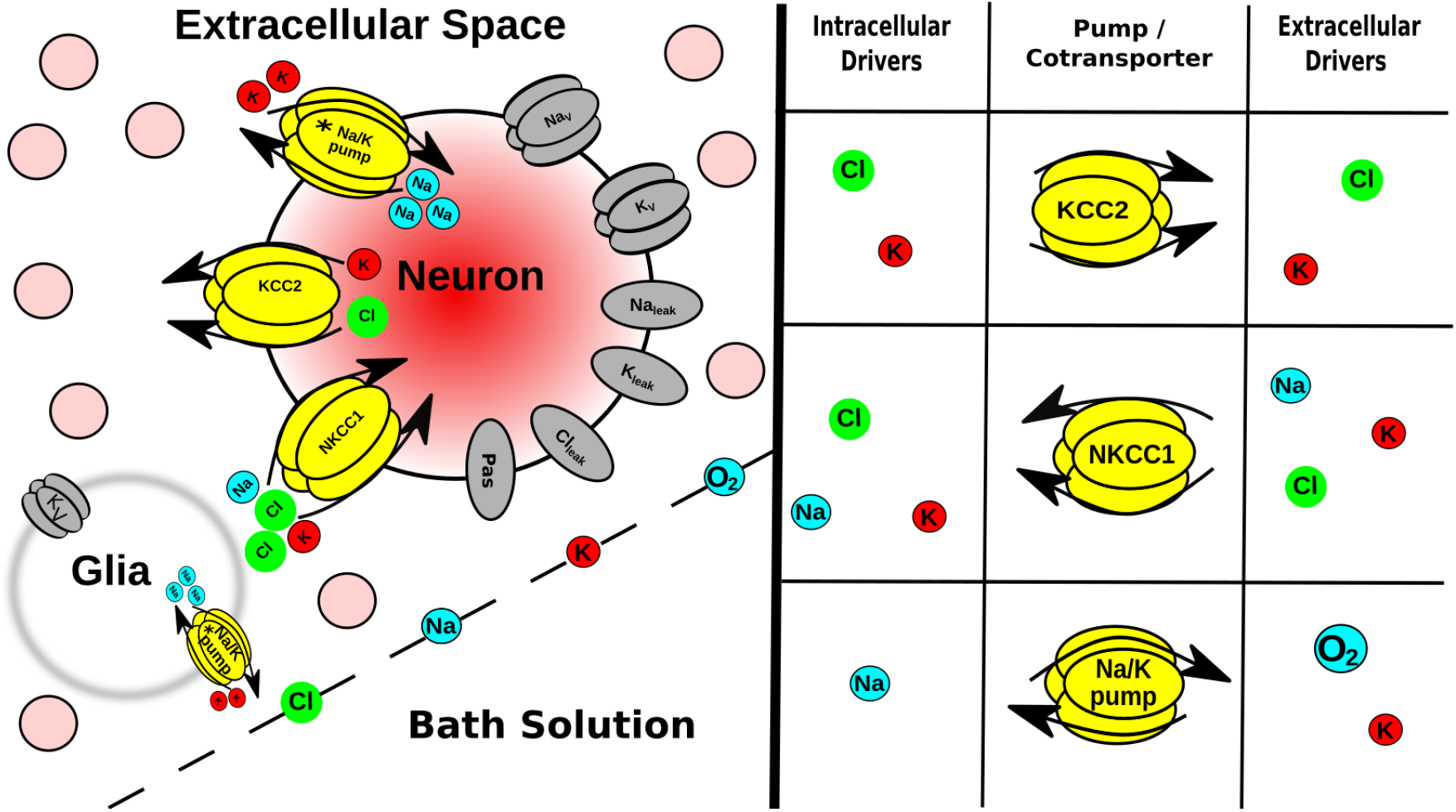
Multiscale model expanded. **Tissue scale**: A few of the 36 · 10^3^ neurons (pink circles) embedded in the ECS of a brain slice submerged in a bath solution where ion and O_2_ concentrations were held constant. Glia are not explicitly modeled, but instead were represented as a field of sinks in every ECS voxel. **Cell scale**: Each neuron had ion channels, 2 co-exchangers; Na^+^/K^+^ pump (asterisk indicates ATP/O_2_ dependence) Ions were well-mixed within each neuron (no intracellular diffusion). **Protein scale**: Table (right) indicates species that control the activity of the intrinsic mechanisms in neurons and in glial-field. **Ion scale**: Ions diffused between ECS voxels by Fick’s law using diffusion coefficients in Table 1.

## Materials and methods

We developed a tissue-scale model of SD in slices by extending the framework developed by Wei *et al*. 2014 [Wei et al., 2014] from a single cell in its local micro-environment to 36,000 cells (baseline) embedded in an ECS. We used the NEURON simulator and its extracellular reaction-diffusion framework (RxD-ECS) to simulate the electrophysiology of those neurons, the exchange of ions between them and the ECS, diffusion of ions and O_2_ through the slice, and exchange of ions and O_2_ between the slice and the bath solution in which it was submerged [Newton et al., 2018]. Our model is not specific to any particular brain area, as we aimed to reproduce general properties applicable to different brain regions and to different pathophysiologies.

**Table 1:** Diffusion coefficients and baseline concentrations for ions and O_2_ in perfused slice [Haynes, 2014, Samson et al., 2003, Wei et al., 2014]

| Species | D ($\cdot 10^{-5}$ cm<sup>2</sup>/s) | Intracellular Concentration (mM) | ECS Concentration (mM) |
| --- | --- | --- | --- |
| $\text{K}^+$ | 2.62 | 140.0 | 3.5 |
| $\text{Na}^+$ | 1.78 | 18.0 | 144.0 |
| $\text{Cl}^-$ | 2.10 | 6.0 | 130.0 |
| $\text{O}_2$ | 3.30 | 0.1 | 0.1 |

## Model

Model neurons were all point-neurons (1 compartment) which included voltage-gated K^+^ and Na^+^ channels; Na^+^, K^+^, and Cl^−^ leak channels; K^+^-Cl^−^ (KCC2) [Payne et al., 2003] and Na^+^-K^+^-Cl^−^ (NKCC1) cotransporters; and Na^+^/K^+^ pumps (Fig. 1), ported from Wei *et al*. (2014) [Wei et al., 2014]. In order for cells to balance a steady-state resting membrane potential in a slice with dynamic ion and O_2_ concentrations, we added a passive electrical conductance with reversal potential E_rev_= −70 mV and conductance *g* = 0.1 mS/cm^2^. Neurons were closed-ended cylinders (no flux from ends). The Na^+^/K^+^ pump activity was dependent on the local concentration of O_2_ in the slice, rather than ATP, and Na^+^/K^+^ pumps were the only consumers of O_2_ in the model [Wei et al., 2014]. ATP reserve in rat cells is estimated to be ∼2.6mM [Veech et al., 1979]. A single-cell simulation using a Michaelis-Menten approximation for ATP-dependence on O_2_ [Noske et al., 2010] demonstrated that [K^+^]_*ECS*_=15 mM increased pump activity by ∼1.8*×* which would reduce reserves to below 1% of within 2 s under hypoxic conditions.

To explore the effects of surface area to volume (S:V) ratio, we used RxD-ECS to independently define a neural surface entirely separated from its volume – hence NOT following the overall geometry of the structure. This is possible since we used the concept of *fractional volumes*, rather than providing actual volume-occupying structures [McDougal et al., 2013]. Neuronal volume fraction *β*_nrn_ is defined as the ratio of total neuronal volume (Vol_nrn_) to total tissue volume: 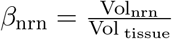 (compare with *α*_*ECS*_ which is 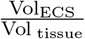)[Rice and Russo-Menna, 1998]. Given a chosen tissue volume, *β*_nrn_, and a total number of neurons *N*_nrn_ (based on cell density times Vol _tissue_), the volume of a single neuron vol_nrn_ is:

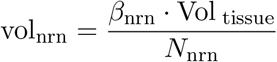

(note case: vol_nrn_ for single cell; Vol_nrn_ for cumulative neuronal volume). In NEURON, neural compartments are cylinders, omitting the ends. In the present case, our point-neurons are each a single cylinder, defined by length L and diameter d. Setting *L* = *d* for simplicity, the surface area is defined as *S*_nrn_ = *π · d*^2^. The associated volume, calculated for this cylinder, was not used. We therefore used RxD-ECS’s FractionalVolume class to scale the volume of a cell with the desired S:V to vol_nrn_, while the cell retains its original *S*_nrn_[McDougal et al., 2013].

To establish a biologically realistic range for S:V, we analyzed morphological reconstructions of neurons from neuromorpho.org. We used results for cells with intact soma and dendritic reconstructions in 3 dimensions from animals older than two weeks. In rat neocortex, average S:V was 3.4 *±* 1.2 *μ*m^−1^ for both pyramidal cells (n=96) and interneurons (n=108) [Radman et al., 2009, Boudewijns et al., 2013, Larkum et al., 2004, Vetter et al., 2001, Meyer et al., 2010, Kubota, 2014]. Higher S:V was grossly associated with larger dendritic trees; S:V scales inversely with diameter in cylindrical structures (excluding ends): 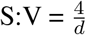. For our baseline simulations, we used S:V = 3.0 *μ*m^−1^; *β*_*nrn*_ = 0.24; neuronal density = 90*·*10^3^ neurons/mm^3^, typical of neocortex [Rice and Russo-Menna, 1998, Keller et al., 2018].

Simulated slices were 1mm x 1mm and ranged in thickness 100 – 800 *μ*m with 45*·*10^3^ – 120*·*10^3^ neurons/mm^3^. The baseline simulation was 400 *μ*m slice with 90*·*10^3^ neurons/mm^3^ (36*·*10^3^ neurons in total). Neurons were situated randomly throughout extracellular space (ECS) with diffusion of 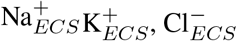, and O_2_ with diffusion coefficients (D) given in Table 1. Extracellular volume fraction (*α*_*ECS*_: ratio of extracellular space volume to total tissue volume) and tortuosity (*λ*_*ECS*_: hindrance to diffusion in tissue compared to free medium) were the same for all ions. This results in a lower effective diffusion coefficient (D*) for ions but not O_2_, which diffused through the slice unhindered. Diffusion through the ECS was calculated with a voxel size of 25 *μ*m x 25 *μ*m x 25 *μ*m. Simulated slices were submerged in simulated bath solution where ion and O_2_ concentrations were equivalent to those estimated for ECS (Table 1, [Cressman et al., 2009, Wei et al., 2014]) with Dirichlet boundary conditions.

In the perfused slice, [O_2_] = 0.1 mM (corresponding to a bath solution aerated with 95% O_2_:5% CO_2_ [Wei et al., 2014]), *α*_*ECS*_ = 0.2, and *λ*_*ECS*_ = 1.6 [Syková and Nicholson, 2008]. We modeled the effects of propionate, which “primed” the tissue for SD, by reducing *α*_*ECS*_ to 0.12 and total Cl^−^ content in the slice by 50%, but keeping [O_2_] = 0.1 mM and *λ*_*ECS*_ = 1.6 [Hrabe and Hrabetova, 2019]. In both cases, SD was initiated by elevating initial [K^+^ ]_ECS_ within a spherical bolus at the center of the slice at t=0. Baseline simulations were run with K^+^ bolus with radius = 100 *μ*m; [K^+^ ]_ECS_ = 70 mM. Hypoxia can also induce a depolarization, termed by some hypoxic SD-like depolarization (HSD), without applying a K^+^ bolus [Pérez-Pinzón et al., 1995, Aitken et al., 1998]. Experimentally, switching the gas to 95% N_2_:5% CO_2_ leads to HSD within minutes, but immediately preceding the depolarization, Perez-Pinzon et al. (1995) identified a period where *α*_*ECS*_ is reduced and *λ*_*ECS*_ is increased which they termed the *preanoxic depolarization phase*. To simulate these conditions, we reduced *α*_*ECS*_ to 0.13, increased *λ*_*ECS*_ to 1.8 [Pérez-Pinzón et al., 1995], reduced [O_2_ ] in the slice to 0.01 mM, and increased K^+^ to 15 mM in a 100 *μ*m radius sphere in the center of the slice, providing a nidus for depolarization initiation. In both Perez-Pinzon et al. (1995) and Aitken et al. (1998), the slice was reperfused immediately after detecting the depolarization, so we set the boundary conditions such that [O_2_ ] in the bath was 0.1 mM (full bath oxygenation). We also simulated a scenario in which a 70 mM bolus of K^+^ was applied to the slice during the preanoxic depolarization phase but before the HSD occurs, and the slice was not reperfused in order to make direct comparisons between SD in an hypoxic and SD in a perfused slice. This could be accomplished experimentally by continuously monitoring ECS properties to determine when to apply the K^+^ bolus.

Studies have shown that *α*_*ECS*_ changes dynamically over the course of SD [Mazel et al., 2002, Hrabe and Hrabetova, 2019]. Since the biophysics of ECS changes during SD have not been elucidated at the time scale of our simulations, we incorporated these changes phenomenologically for a subset of simulations, reducing *α*_ECS_ to 0.05-0.1 during passage of the SD [Mazel et al., 2002]. In each ECS voxel:

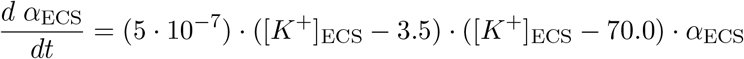

This model of dynamic *α*_*ECS*_ only accounts for the drop in *α*_*ECS*_ during SD, not its recovery after SD, which occurs on the time-scale of minutes [Mazel et al., 2002, Hrabe and Hrabetova, 2019].

Glia were modeled by a background voltage-gated K^+^ current and Na^+^/K^+^ pump in each ECS voxel [Cressman et al., 2009, Øyehaug et al., 2012, Wei et al., 2014], rather than as individual cells.

## Simulations

SD was initiated with K^+^ bolus with radius = 100 *μ*m; [K^+^ ]_ECS_ = 70 mM, unless noted otherwise. To follow the position of SD over time, we tracked the position of the K^+^ wavefront, defined as the furthest location where [K^+^ ]_ECS_ exceeded 15 mM, and the positions of spiking neurons. Propagation speed was indicated as 0 if the K^+^ wave didn’t propagate. Most simulations ran for 10 seconds, which was sufficient for SD to propagate throughout the entire slice.

In the course of this study, we ran over 600 simulations covering a range of slice sizes, cell densities, and durations on a number of different architectures. Simulating a 1 mm x 1 mm x 400 *μ*m slice with a cell density of 90k neurons/mm^3^ (36*·*10^3^ neurons) for 10 s of simulation-time took approximately 12.5 hours on a parallel Linux system using 48 nodes on a 2.40 GHz Intel Xeon E5-4610 CPU. Incorporating dynamic *α*_*ECS*_ into the same model on the same machine increased simulation time to approximately 18 hours. Due to these computational limitations we restricted out simulations to 10 sec which meant that we could not continue to the termination phase of SD which would require considerably greater temporal and spatial scales. Simulations were run using Neuroscience Gateway [Sivagnanam et al., 2013], Google Cloud Platform, and the on-site high performance computer at SUNY Downstate Health Sciences University.

## Results

In our model of an O_2_-perfused slice, a small bolus of elevated K^+^ (70 mM, 100 *μ*m radius) initiated a propagating K^+^ wave with associated spreading depolarization (SD) producing neuronal spiking (Fig. 2). The K^+^ wave travelled radially outward from the bolus in 3 dimensions towards the edges of the slice at 2.3 mm/min, comparable to optical and electrophysiological measurements of SD propagation velocity in brain slices [Aitken et al., 1998, Joshi and Andrew, 2001, Hrabe and Hrabetova, 2019]. Within the K^+^ bolus, most cells fired a single spike and went immediately into depolarization block. Outside the K^+^ bolus, cells fired a 200-900 ms burst of action potentials as [K^+^ ]_ECS_ increased around them. During the course of the SD-associated burst, instantaneous firing rates increased to as high as 250 Hz with decreasing spike heights during the burst, comparable to experimental observations [Devin Brisson et al., 2013, Lemaire et al., 2021]. Cells then remained in depolarization block for the remainder of the 10 s measured (see *Methods* for computational limitations) [Devin Brisson et al., 2013, Andrew, 2016]. Spreading *depolarization*, seen intracellularly, produced Na^+^ channel inactivation and prevented further spiking. Absence of spiking would be seen extracellularly as spreading *depression* – a silent region in the slice. The K^+^ wave and SD were coincident in time and space, with spreading depression following closely behind; we primarily followed the K^+^ wave since this was easiest to localize across conditions.

**Figure 2:**
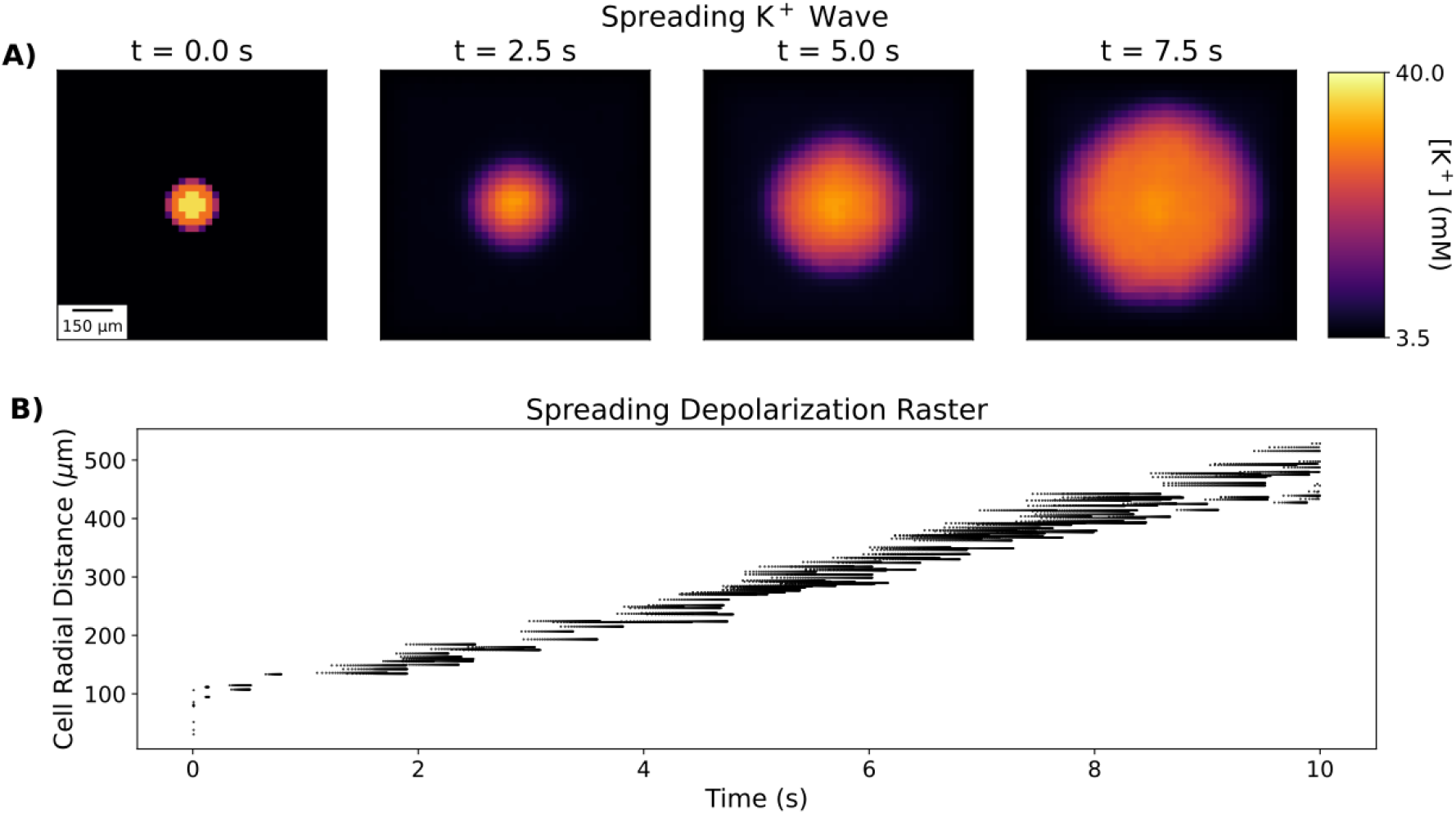
A small bolus of applied K^+^ initiates spreading K^+^ and depolarization waves in perfused slice. A. [K^+^ ]_ECS_ averaged across slice depth (400 *μ*m) at 4 time points during SD. B. Spike raster plot of 250 randomly-selected neurons (out of 36 · 10^3^) during SD. Cells are ordered on y-axis by their radial distance from the center of the K^+^ bolus. Blank area under spikes represents region of spreading depression. Baseline values: [O_2_] = 0.1 mM; *α*_*ECS*_ = 0.2; *λ*_*ECS*_ = 1.6; [Cl^−^]_*ECS*_ = 130.0 mM; [Cl^−^]_*i*_ = 6.0 mM.

SD in perfused slices was an all-or-none process; it could only be initiated above a certain threshold measured either in concentration – [K^+^ ]_ECS_ ≥ 20 mM (bolus diameter 200*μ*m), or in bolus diameter – diameter ≥ 100*μ*m ([K^+^ ]_ECS_ =70 mM) (Fig. 3). Beyond these thresholds, different K^+^ bolus concentrations and diameter had only a minimal effect on wave speed.

**Figure 3:**
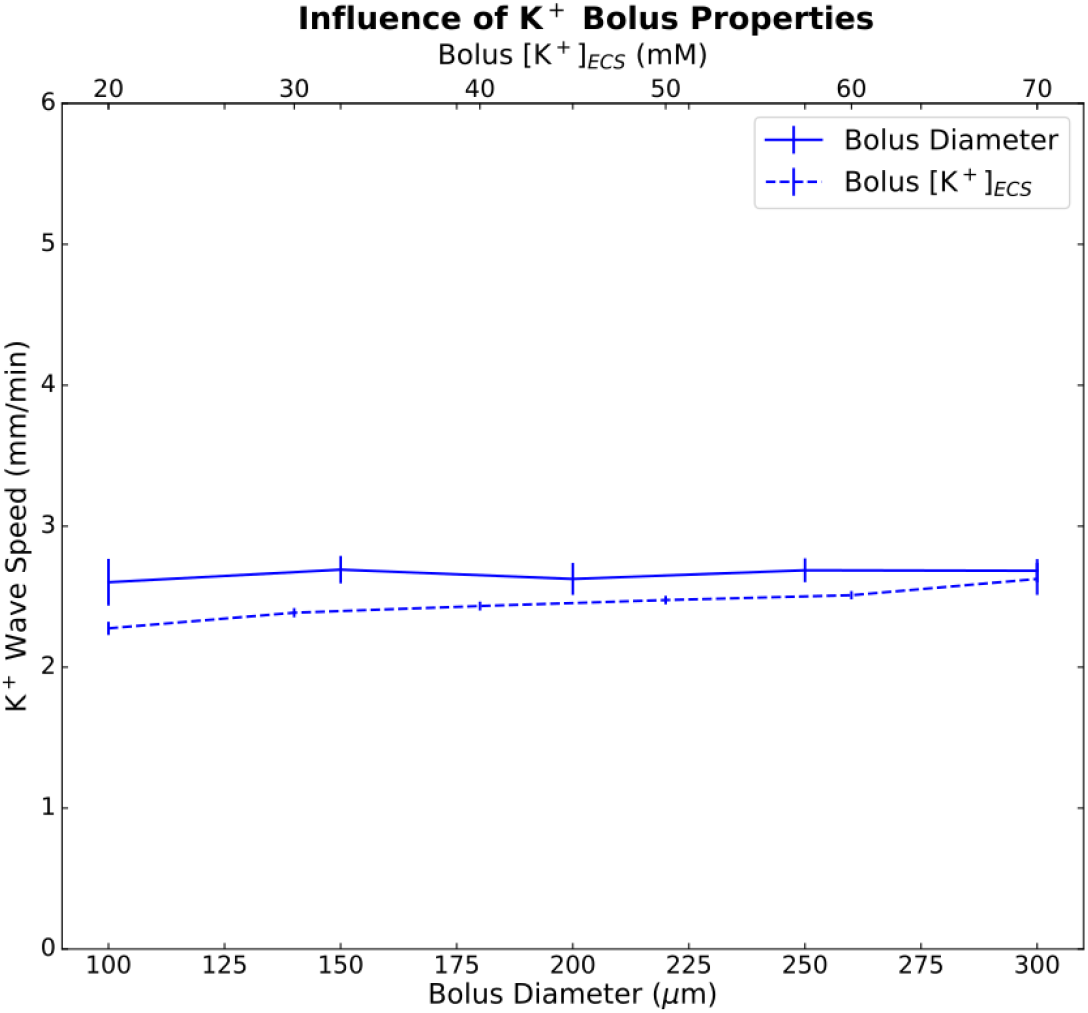
Diameter (bottom x-axis) and concentration (top x-axis) of the K^+^ bolus had minor effects on K^+^ wave speed. Mean and standard deviations (*n* = 5) for K^+^ wave speed versus bolus diameter (solid line; bolus [K^+^ ]_ECS_ =70 mM; 5 random cell position initializations) and versus bolus concentration (dashed line; diameter=200 *μ*m).

Underpinning SD was a wave of pronounced imbalance of transmembrane ion concentrations (Fig. 4, Supporting Movie 4-1). Excess K^+^ is briefly eliminated from the ECS via neural and glial homeostatic mechanisms. Once the K^+^ wave arrived however, the cells dumped large quantities of K^+^ into the ECS due to the burst and subsequent prolonged depolarization (note rapid depletion of [K^+^ ]_i_ – Fig. 4B). The Na^+^/K^+^ pump activity which contributed to K^+^ elimination from the ECS in the core of the slice created a high demand for O_2_, exceeding the rate at which it could diffuse in from the bath. This resulted in much of the tissue becoming hypoxic before the arrival of the K^+^ wave (compare rapid fall-off of O_2_ in Fig. 4E to much slower rise of extracellular K^+^ in Fig. 4F). The rapid spread of O_2_ deficit explains the total pump failure at intermediate locations in the slice. There were small upward deflections in the first 3 traces in Fig. 4B reflecting homeostatic inward pumping. There was no upward deflection in the other 3 traces – O_2_ has disappeared before the K^+^ wave arrives. The preservation of pumping in the final, most peripheral, trace of Fig. 4B is due to this measurement being at the edge of the slice, neighboring the O_2_ source of the bath. In reality, neuronal ATP reserves will maintain pumping for a limited duration in the absence of O_2_, but reserves will quickly run out due to the high metabolic demand imposed on the pumps by large shifts in K^+^ and Na^+^concentrations. Once [K^+^ ]_ECS_ reached ∼14 mM, cellular homeostatic mechanisms totally broke down. Changes in intra- and extracellular Na^+^concentrations were actually larger than the shifts in K^+^ accompanying SD, and a wave of extracellular Na^+^deficit traveled along with the K^+^ wave (Fig. 4C,G). Shifts in Cl^−^concentrations proceeded through the slice slightly more slowly and were less pronounced (Fig. 4D,H).

**Figure 4:**
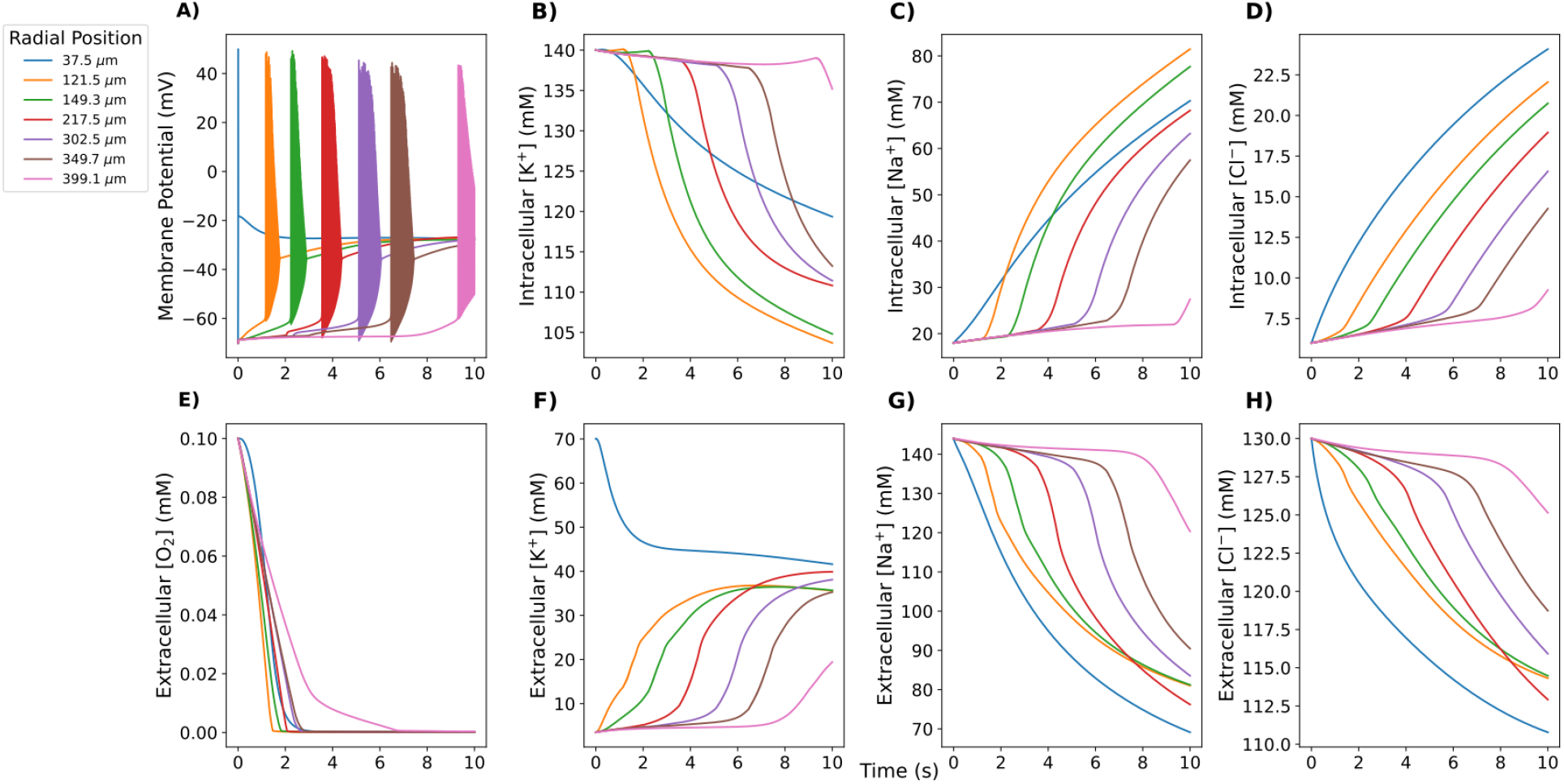
Concentrations at 7 radial locations measured during SD in perfused 400 *μ*m slice) A. Cell within the K^+^ bolus (37.5 *μ*m) produced a single spike; cells farther out fired a burst. Cells remained in depolarization block for the remainder of the simulation (10 s). B-D. Intracellular ion concentrations. E-H. Extracellular O_2_ and ion concentrations in neighboring ECS voxels. Supporting Movie 4-1 shows extracellular ion and O_2_ concentrations across the slice, as well as neuronal spiking (white dots) from 250 neurons during the course of SD (most easily seen with slowed playback).

SD was facilitated by incorporating the effects of hypoxia or propionate treatment on the slice. Hypoxic SD-like depolarization (HSD, caused by hypoxia, rather than introducing a K^+^ bolus) propagated similarly to SD in perfused slice, as shown experimentally [Aitken et al., 1998] (Fig. 5). Perez-Pinzon et al. (1995) identified a period immediately preceding the HSD they termed a *preanoxic depolarization phase*, with reduced *α*_*ECS*_and increased *λ*_*ECS*_. We modeled HSD by incorporating these hypoxia effects, using a small region of slightly elevated K^+^ (100 *μ*m radius, 15 mM) to provide a nidus for HSD initiation. The spatiotemporal distribution of neuronal spiking was similar to that of the control (perfused) slice, as seen experimentally. (Fig. 5B, Supporting Movie 5-1). However, the K^+^ wave during HSD was faster (3.7 mm/min) than during SD in perfused slice (2.3 mm/min) (Fig. 5A), We also simulated standard K^+^-initiated SD in hypoxic slices by applying a 70 mM K^+^ bolus (as with SD in perfused slice) to a slice during this preanoxic depolarization phase [Pérez-Pinzón et al., 1995], resulting in K^+^ wave speed of 3.4 mm/min. Simulating propionate application (decreased *α*_*ECS*_=0.12; halving [Cl^−^]_*i*_ and [Cl^−^]_*ECS*_ [Tao et al., 2002, Hrabe and Hrabetova, 2019]), increased K^+^ wave speed to 4.8 mm/min (Fig. 5A). Comparable speedups were also observed in the depolarization waves (Fig. 5B). Since these manipulations included combined changes to [O_2_], Cl^−^, *α*_*ECS*_, and/or *λ*_*ECS*_, we investigated their individual contributions over the relevant ranges. Since both hypoxia and propionate decrease *α*_*ECS*_, we also tested increasing *α*_*ECS*_ to as high as 0.42, which has been observed when making the ECS artificially hypertonic [Kume-Kick et al., 2002]. *α*_*ECS*_ had the largest influence on propagation, changing K^+^ wave speed by *>*2 mm/min over the range tested, while K^+^ wave speed changed by *<*0.5 mm/min for the ranges of [O_2_], *λ*_*ECS*_, and [Cl^−^] values tested (Fig. 5C).

**Figure 5:**
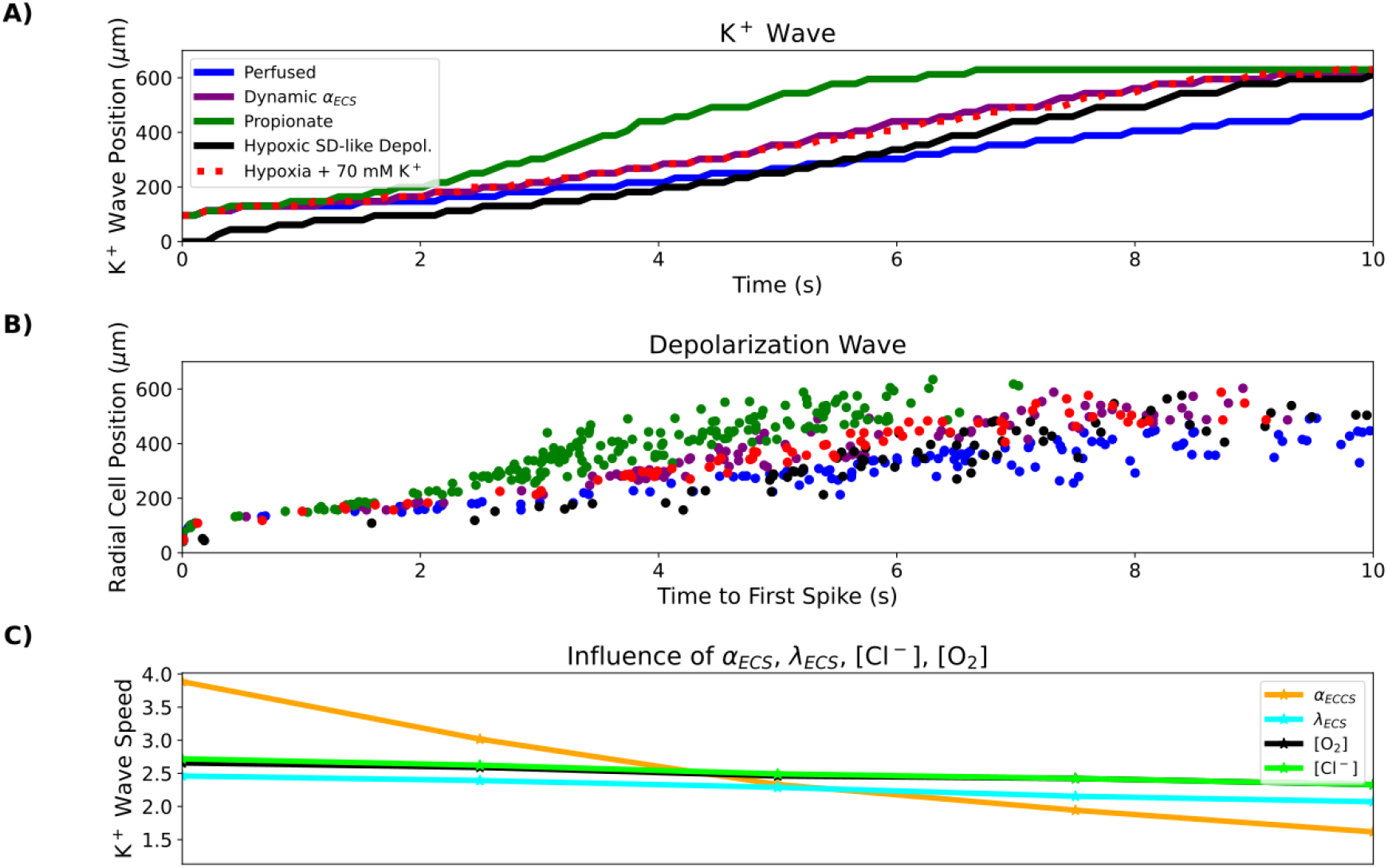
Hypoxia, propionate, and dynamic ECS increased SD speed principally through *α*_ECS_ reduction. A. Radial K^+^ wave position over time during SD in perfused, hypoxic, HSD, propionate conditions, and with dynamic changes in *α*_**ECS**_. (Supporting Movie 5-1). Hypoxia, propionate, and dynamic changes in *α*_ECS_ facilitated propagation. B. Radial position of SD wave represented by time to first spike in 126 selected cells at different distances from center. C. K^+^ wave speeds with individual parameter changes (Fig. 2). *α*_ECS_ had the greatest impact on SD speed over physiologically plausible range. (x-axis ranges: [O_2_]=0.01-0.1 mM; *α*_*ECS*_=0.07-0.42; *λ*_*ECS*_=1.4-2.0; [Cl^−^]_*ECS*_:[Cl^−^]_*i*_=3.0:65.0-6.0:130.0 mM)

Experimental studies have demonstrated dynamic changes in *α*_*ECS*_ occurring during SD, with *α*_*ECS*_ dropping to as low as 0.05 at the peak of the depolarization [Mazel et al., 2002, Hrabe and Hrabetova, 2019]. Given the strong influence of constant *α*_*ECS*_ on SD propagation (Fig. 5C - orange line), we also explored the influence of dynamic *α*_*ECS*_ (Fig. 5A - purple line). Dynamically decreasing *α*_*ECS*_ was modeled as a local function of increasing [K^+^ ]_ECS_ (see *Methods*), such that *α*_*ECS*_ dropped to 0.06 in the wake of SD, within the experimentally-observed range of 0.05 - 0.1 [Mazel et al., 2002]. Incorporating a dynamic ECS increased the speed of SD propagation in perfused slice from 2.3 mm/min to 3.3 mm/min. The stereotyped characteristics of neuronal firing patterns during depolarization wave remained unchanged by dynamic changes in *α*_*ECS*_.

Different brain areas have different cell densities and their neurons have different morphological characteristics. We manipulated our generic model so as to explore 3 properties of neural tissue organization and shape: neuronal surface area to volume ratio (S:V); fraction of tissue volume occupied by neurons (*β*_nrn_); and cell density (number of neurons per mm^3^) (Fig. 6). Neuronal S:V varies across cell types, brain regions, and species. Examination of representative morphologies showed that S:V values are generally in a range of 2–10 *μ*m^−1^ (see *Methods*), with neocortical principal cell S:V of 3.4 *±* 1.2 *μ*m^−1^ (n=96) significantly greater than brainstem principal cell S:V of 2.2 *±* 1.2 *μ*m^−1^ (n=74; p*<*0.001, Mann-Whitney U test) [Williams et al., 2019, Ros et al., 2010, Núñez-Abades et al., 1994, Raslan et al., 2014, Radman et al., 2009, Boudewijns et al., 2013, Larkum et al., 2004, Vetter et al., 2001, Meyer et al., 2010]. Neuronal volume fraction *β*_nrn_ may differ with different brain areas, and will differ under the pathological condition of cytotoxic edema. Cell density varies across different neural areas.

**Figure 6:**
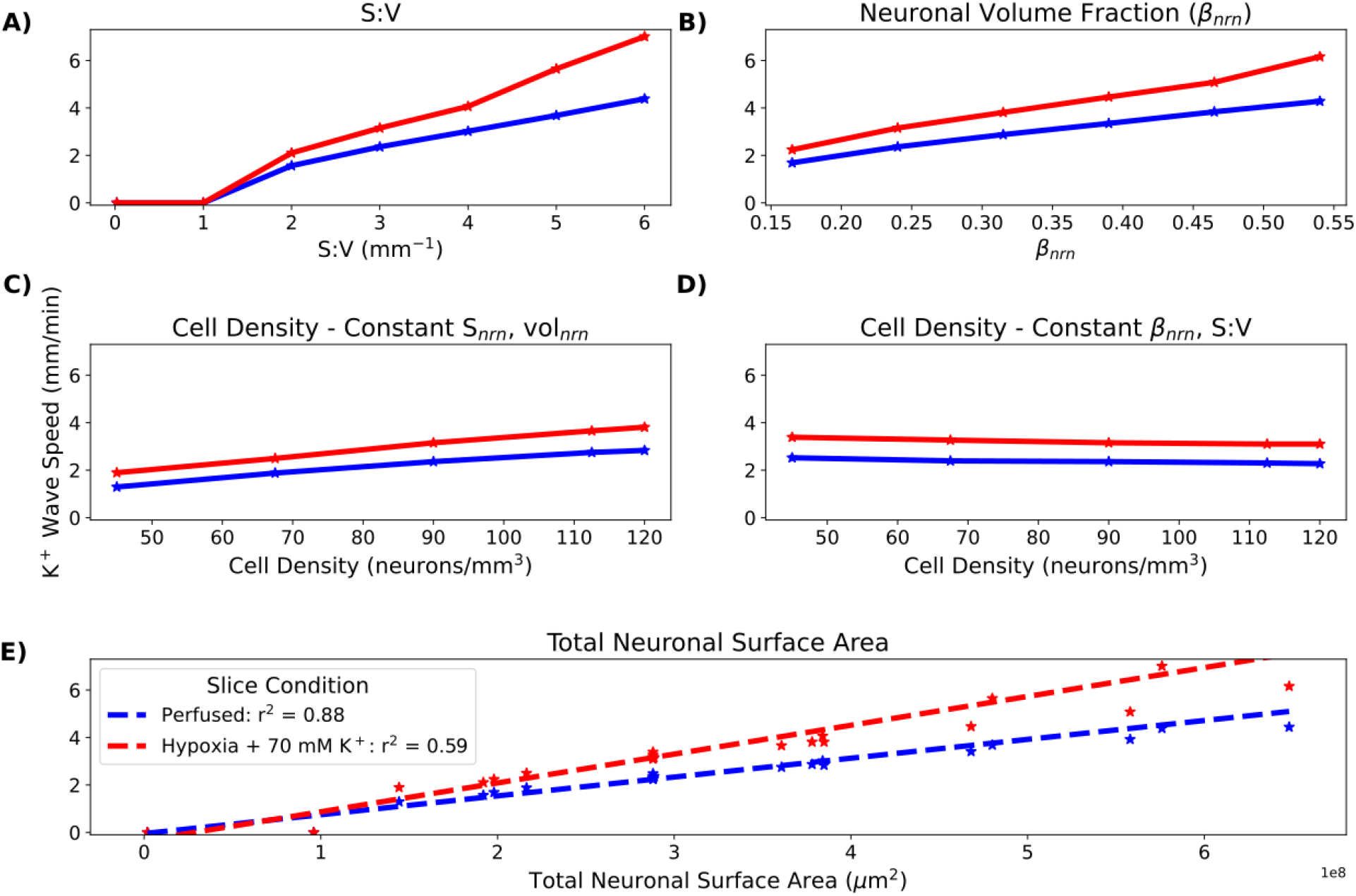
K^+^ wave propagation speed proportional to total neuronal surface area in slice. SD initiated in perfused and hypoxic slices (no reperfusion) by introducing a 100 *μ*m radius central 70 mM spherical K^+^ bolus. Effects of A. varying S:V of each cell while maintaining a constant *β*_*nrn*_; B. varying *β*_*nrn*_ while keeping S:V constant; thus allowing S_*nrn*_ to vary; C. varying cell density while keeping constant S_*nrn*_ and vol_*nrn*_; thus allowing *β*_*nrn*_ to vary; D. varying cell density while keeping *β*_*nrn*_ and S:V constant; thus allowing S_*nrn*_ and vol_*nrn*_ to vary. E. Pooled results: K^+^ wave speed increased linearly with total neuronal surface area in both perfused and hypoxic slices. Hypoxia increased wave speed across conditions (0 speed indicates no SD).

Realistic (*>*1 *μ*m^−1^) S:V was necessary for initiating SD (Fig. 6A) – SD could not be initiated using the actual 3D geometry of single-cylinder point-neurons with diameter and height selected to produce baseline *β*_nrn_ of 0.24 (S:V = 0.02 *μ*m^−1^ with 90k neurons/mm^3^, perfused or hypoxic). Above this threshold, K^+^ wave speed increased with S:V. K^+^ wave speed also increased with increased *β*_nrn_ while keeping S:V and cell density (number of neurons per mm^3^) constant (Fig. 6B). Cell density effects were less marked, whether through keeping surface area and volume constant (Fig. 6C), or keeping *β*_nrn_ and S:V constant (Fig. 6D). In all cases, change with altered parameters was more pronounced in the hypoxic slice. Pooled together, we found a near linear relationship of K^+^ wave speed with total neuronal surface area

Slice thickness (100–800 *μ*m) influenced SD by altering the ability of O_2_ to penetrate to the tissue core (Fig. 7). SD could not be initiated in 100 *μ*m perfused slice – SD was not sustainable with full O_2_ availability, but it was initiated in an hypoxic slice of the same thickness. With increasing thickness, an increasingly hypoxic core (despite O_2_ perfusion of the bath) allowed K^+^ wave speed to increase from 1.2-2.1 mm/min over 200-400 *μ*m thickness (Fig. 7A). Above 400 *μ*m there was no increased speed with increased thickness. Similar patterns were observed in hypoxic slices.

**Figure 7:**
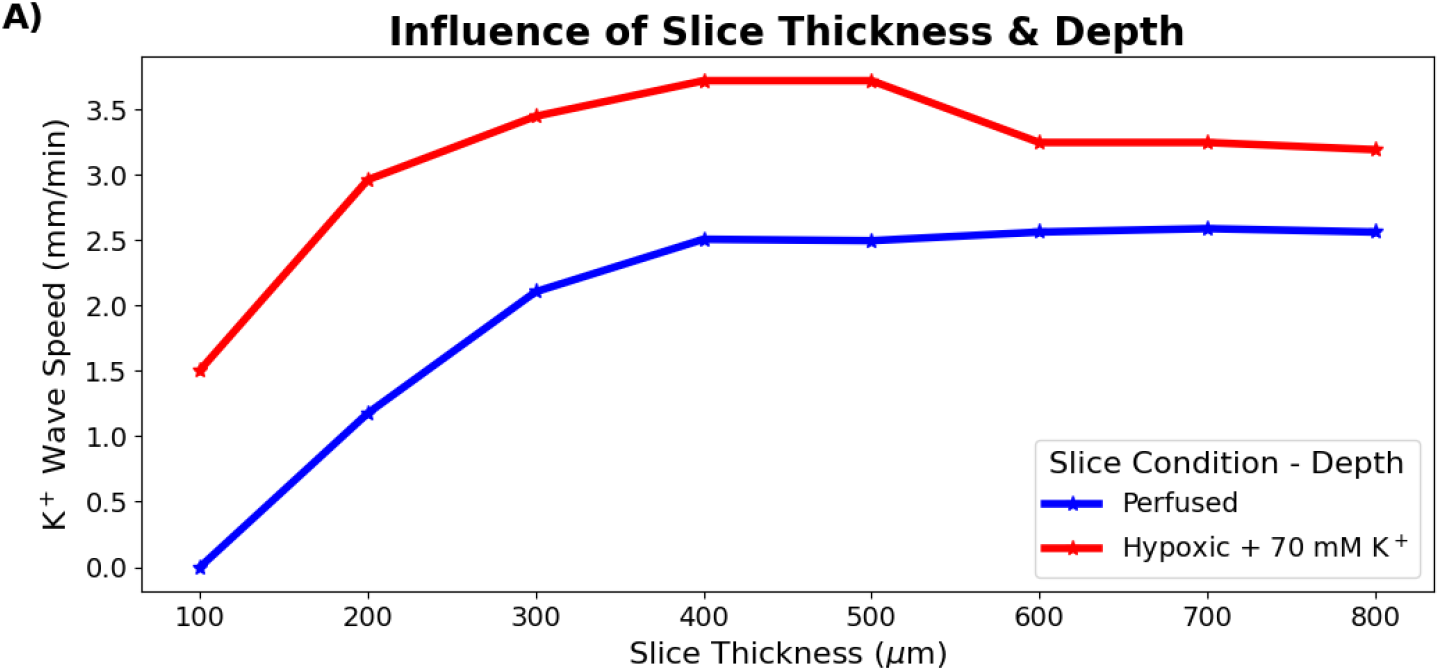
Slice thickness effects on SD propagation. A. K^+^ wave speed during SD in perfused and hypoxic slices of various thicknesses. SD could not be initiated in very thin (100 *μ*m) slices when perfused but could in hypoxic slices. For perfused slices, K^+^ wave speed increased with slice thickness between 200-400 *μ*m then saturated for slices of greater thickness. A similar pattern was observed in the hypoxic slices with consistently faster K^+^ wave speeds compared to in perfused slice.

We also observed depth-dependent differences in propagation of the SD within a slice (Fig. 8). A wave of high K^+^ propagated through the core of the slice (*±*50*μ*m from the center, Fig. 8A), and neurons there exhibited the membrane dynamics of SD: a burst of spikes followed by depolarization block lasting the remainder of the simulation (Fig. 8C). The K^+^ wave reached the periphery of the slice (within 50 *μ*m of bath) but in lower concentration (Fig. 8B), and neurons there were resistant to SD due to the high availability of O_2_ from the bath (Fig. 8D). Instead, neurons in the periphery of the slice only underwent a modest depolarization and fired regularly at 10-70 Hz, without the bursting followed by depolarization block characteristic of SD.

**Figure 8:**
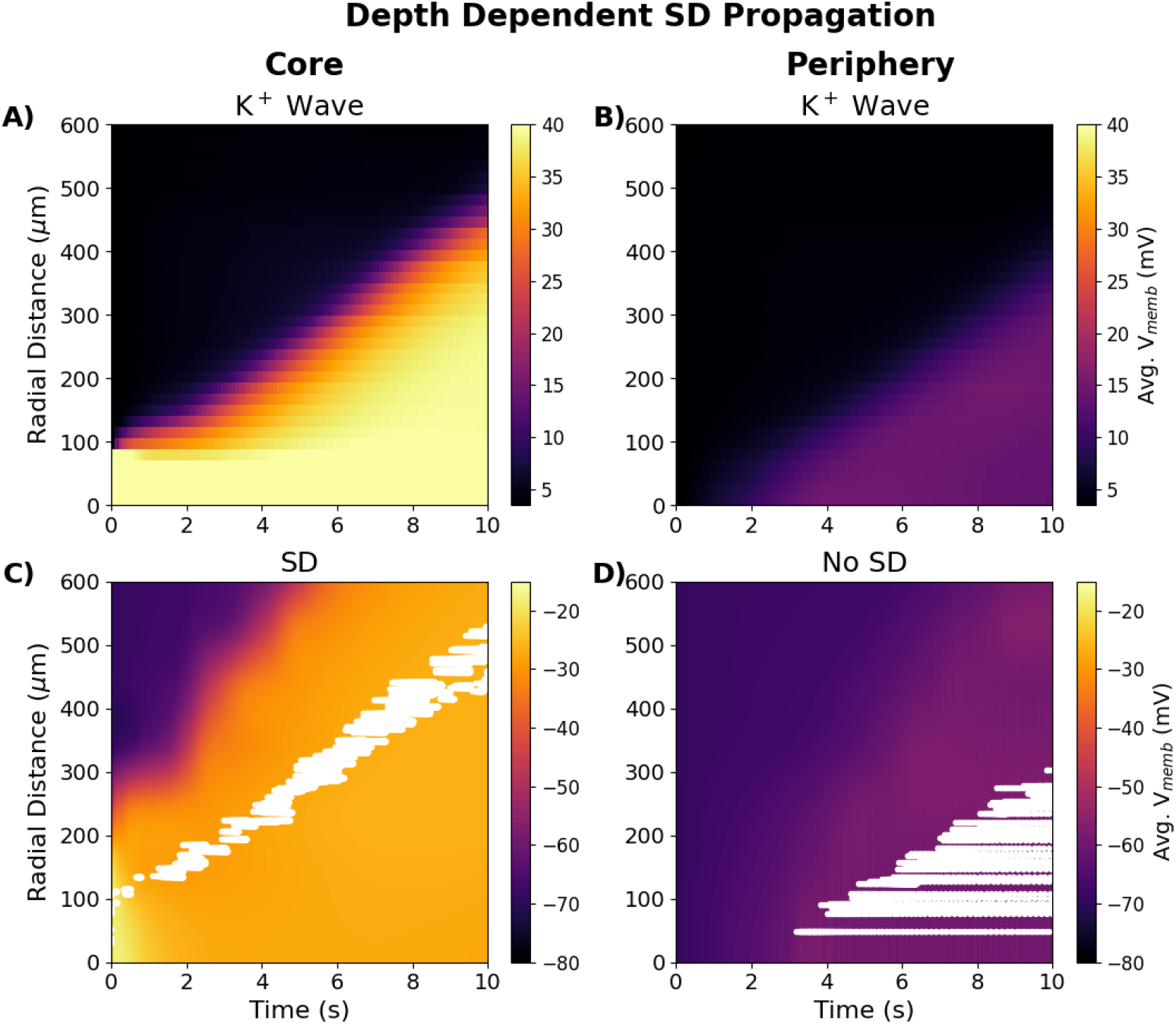
Depth-dependence of SD propagation in 400 *μ*m thick perfused slice. A. Spread of K^+^ wave through slice core (*±* 50*μ*m from center). B. Wave of mildly elevated K^+^ reached the periphery (within 50 *μ*m of bath) from the central bolus. C. Spread of SD through the slice core. Voltages in slice core (color map) with spike rasters for 120 cell subset overlaid in white. Neurons in the core showed typical SD voltage dynamics (bursting followed by depolarization block). D. Voltages and raster at periphery – cells show regular spiking patterns at 10-70 Hz. (V_*memb*_ averaged across all cells in 25*μ*m*×* 25*μ*m*×* 100*μ*m voxels at center or periphery)

## Discussion

Our model reproduced a number of the properties of SD observed in brain slices (Fig. 2,4). Slice models which most resembled cortical gray matter (high cell density, high neuronal S:V) showed SD speeds and neuronal firing patterns comparable to *in vitro* optical and electrophysiological measurements [Aitken et al., 1998, Hrabe and Hrabetova, 2019, Devin Brisson et al., 2013, Lemaire et al., 2021]. The all-or-none nature of SD initiation, as well as the observed bolus [K^+^ ] threshold of ∼20 mM, was also in agreement with experiments in brain slices [Andrew et al., 2017].

Our simulations identify a pathological cycle whereby SD induces, and is also exacerbated by, hypoxia in slice. *In vivo*, SD-associated metabolic demand for O_2_ can exceed supply, resulting in tissue hypoxia [Takano et al., 2007, Piilgaard and Lauritzen, 2009]. However, *in vivo* experiments showed hypoxia following SD, rather than preceding it as predicted by our *in vitro* simulations. *In vivo*, O_2_ is supplied by the vasculature, and increased neural activity during SD may lead to increased blood supply and tissue oxygenation [Balança et al., 2017], while *in vitro*, O_2_ is only supplied by the bath which is effectively unchanging. While K^+^ was slowly spreading outward across the slice, O_2_ spread inward rapidly, following the gradient caused by O_2_ depletion from overworked pumps, but was quickly consumed (compare rapid fall-off of O_2_ in Fig. 4E to much slower fall-off of [K^+^ ]_i_ in 4B). Depending on the distance to the bath and to the inciting bolus, intermediate locations in the slice suffered various degrees of pump demand and partial pump failure. The resistance to SD of neurons in the periphery of the slice compared to those in the core (Fig. 8B) was comparable to findings *in vivo*: tissue near capillaries resist SD, while tissue further away is relatively hypoxic and prone to SD [Takano et al., 2007].

By comparing the effects of changing O_2_ availability, total Cl^−^ content, *α*_*ECS*_, and *λ*_*ECS*_ on SD in isolation, we determined that *α*_*ECS*_ influenced SD most strongly (Fig. 5C), accounting for most of the speed-up observed in hypoxic and propionate-treated slices. Our results support the hypothesis that the main mechanism in propionate’s priming for SD is through reducing *α*_*ECS*_ [Hrabe and Hrabetova, 2019], and suggest that the main mechanism for hypoxia speed-up is also reduced *α*_*ECS*_.

We modeled SD in brain slices as a reaction-diffusion system in an unconnected neuronal population. Although we initiated SD by elevating extracellular K^+^ and tracked the position of K^+^ waves, the model itself was agnostic as to the agent of SD propagation. In our simulations of HSD, we saw that the initial elevation of extracellular K^+^ did not immediately initiate a spreading wave of K^+^, but led to delayed spread (Fig. 5A). In our simulations, as in slice experiment, when K^+^ was applied to the tissue or hypoxia was induced, elevation of K^+^ preceded the wavefront of SD-associated neuronal spiking. By contrast, when SD has been experimentally initiated in other ways, depolarization may precede, rather than follow, the K^+^ wave [Somjen, 2001]. Incorporating synaptic connections into the model will provide a saltatory forward activation that will tend to speed up depolarization relative to the K^+^ wave; the balance of these factors will depend strongly on cell excitability, and the density and lengths of connections.

### Model limitations

An irony of this type of study is that we are computer modeling an *in vitro* model of an *in vivo* animal model of human pathophysiology (model of a model of a model). There are necessarily distortions at each step. At the computer modeling level, the major limitations of this study are the limitations that are inherent in all computer modeling -— we necessarily made choices as to what to include and what to leave out. In the current set of simulations, we left out: **1**. all neural connectivity; **2**. types of neurons, including distinction between inhibitory and excitatory neurons; **3**. dendritic morphologies + additional membrane mechanisms; **4**. glia, except as a generalized field; **5**. volume occupying structures (instead using fractional volumes); **6**. intracellular handling of ions and second messengers with effects on pumps and other mechanisms. These are largely correctable limitations that we will gradually begin to address in future versions of the model. Some, however, represent limitations in experimental knowledge which need to be addressed.

Additionally, we purposely designed the model to be generic rather than to reproduce the properties of any particular brain region and species. We feel this allowed us to generalize more readily – *e*.*g*., comparing SD in brainstem vs cortex. Several interdependent tissue properties were treated independently with the benefit of allowing additional investigation by isolating parameters.

A major limitation of the model was the simplification in which Na^+^/K^+^ pumps consumed O_2_ to drive their conductances rather than ATP [Wei et al., 2014]. This was partly justified by Na^+^/K^+^ pumps using 91% of available ATP in the brain under normal physiological conditions [Lennie, 2003]; also, neurons synthesize ATP as needed [Davis, 2020]. Because of ATP reserves in the tissue and the simplification using O_2_ as a proxy for ATP, the spread of hypoxia during SD in perfused slices may be slower than predicted by our simulations, but we still expect it to precede the SD wavefront *in vitro*.

We also note that extracellular Na^+^ concentration shifts following SD were smaller than observed experimentally and intracellular concentration shifts were larger than expected [Somjen, 2001]. This may reflect biophysical mechanisms absent from the model or a lack of appropriate volume changes in ECS and intraneuronal spaces. In particular, we did not model large, intracellular, negatively charged macromolecules that produce Gibbs-Donnan effects which contribute to SD [Lemale et al., 2022].

### Experimentally Testable Predictions

Several of our predictions relate to the effects of manipulations on SD speed. These effects could be most easily assessed electrophysiologically by using a series of extracellular electrodes in slice to note the time of population burst passage and subsequent time of silence (the depression phase).

1. **Slower SD in brainstem slice compared to cortical slice**. Compared to cortex, brainstem has lower cell density, higher *α*_*ECS*_, and low expression of ECS matrix molecules/perineural nets, implying low *λ*_*ECS*_ [Ogawa et al., 1985, Hobohm et al., 1998, Syková and Nicholson, 2008], All of these factors contribute to slower propagation speeds in our model (Fig. 5C, 6) Our analysis of principal cell morphologies from brainstem also suggested a S:V lower than those of neocortical principal cells, another factor contributing to slowing.
2. **Increased SD speed with cytotoxic edema in penumbra after stroke or traumatic brain injury (TBI)**. Cytotoxic edema will increase *β*_*nrn*_, producing speed-up, which will be enhanced in the hypoxic condition (Fig. 6B). We note that some of the fastest SD speeds (∼9 mm/min) have been observed in patients after stroke [Woitzik et al., 2013].
3. **Hypoxia will increase SD speed under multiple conditions**. SD spread faster in hypoxic than in perfused slice. Experimentally, one would assess as follows: 1. remove O_2_ from bath; 2. monitor *α*_*ECS*_ to determine the onset of preanoxic depolarization phase (*α*_*ECS*_ ∼0.099–0.0179, depending on brain region [Pérez-Pinzón et al., 1995]) 3. add K^+^ bolus. The slice should not be reperfused, unlike the procedure for HSD experiments [Pérez-Pinzón et al., 1995, Aitken et al., 1998].
4. **Increasing** *α*_*ECS*_ **will attenuate SD propagation**. This could be assessed using hypertonic saline to increase *α*_*ECS*_ [Kume-Kick et al., 2002]. Hypertonic saline is sometimes used in reducing intracranial pressure after TBI [Oddo et al., 2009, Mangat et al., 2020, Shi et al., 2020, Kamel et al., 2011], and might therefore also reduce SD in these patients.
5. **SD speed will be reduced by anti-epileptic drugs**. We showed here that dynamic changes in *α*_*ECS*_ due to SD itself speeds up the SD wave (Fig. 5A, B). Similar changes in *α*_ECS_ have been seen with ictal phenomena in [Colbourn et al., 2021], allowing us to hypothesize that this may be the linkage between SD and *α*_*ECS*_, presumably mediated through excessive release of neurotransmitters whether classical, peptidergic, or NO.
6. **SD velocity will correlate with neural density, dendritic complexity and total neuronal volume across regions** Measurements of SD in various brain regions and across species can be assessed. Density is determined with counts in Nissl stain. Dendritic complexity increases S:V and can be assessed on traced, biocytin-filled cells with measures such as we performed here. Total neuronal volume can be assessed by measuring ascorbic acid in tissue [Rice and Russo-Menna, 1998]. These effects could also be evaluated in tissue culture or in organoids.
7. **Increasing the diameter or concentration of the K**^+^ **bolus used to initiate SD beyond their thresholds will not change SD speed (Fig. 3)**.
8. **Ease of SD initiation and SD propagation speed will increase with increasing slice thickness due to relatively hypoxic core (Fig. 7)**. SD will be difficult or impossible to initiate in very thin slices.
9. **SD propagates through the core of the slice. Neurons near the surface of a perfused slice will be resistant to SD due to the high availability of O**_2_ **(Fig. 8)**. Extracellular recordings looking for bursting and subsequent depression at different slice depths could be performed to confirm this prediction. However, one may want to avoid measurement directly at the slice surface where neurons will have suffered damage due to slicing and therefore may exhibit additional pathology that could alter SD propagation.

### Future directions

Our model incorporated quantitative data and simpler models from numerous sources and at multiple spatial scales to constitute a unified framework for simulating SD in brain slices. We propose the use of this framework as community tool for researchers in the field to test hypotheses; guide the design of new experiments; and incorporate new physiological, transcriptomic, proteomic, or anatomical data into the framework. The open-source, branchable, versioned GitHub repository can provide a dynamical database for SD simulations or modeling brain slices in general.

## Acknowledgements

This work was funded by NIH NIMH R01MH086638 and NSF Grant Internet2 E-CAS 1904444. We wish to thank Drs. Rena Orman, Mark Stewart, Richard Kollmar, and John Kubie (SUNY Downstate) for useful discussions on this topic. We also with to thank Dr. Michael Hines (Yale) for his support with the NEURON simulator.

## Extended Data

**Extended Data 1 - Simulation code:** Our tissue-scale model of spreading depolarization (SD) in brain slices is available on GitHub. We used the NEURON simulator’s reaction-diffusion framework to implement embed thousands of neurons (based on the the model from Wei et al. 2014) in the extracellular space of a brain slice, which is itself embedded in an bath solution. We initiated SD in the slice by elevating extracellular K+ in a spherical region at the center of the slice. Effects of hypoxia and propionate on the slice were modeled by appropriate changes to the volume fraction and tortuosity of the extracellular space and oxygen/chloride concentrations. Users need to install NEURON, and we recommend using MPI to parallelize simulations.

**Extended Data 2 - Supporting Movie 4-1:** Supporting Movie 4-1 shows extracellular ion and O_2_ concentrations across the slice averaged over depth, as well as neuronal spiking (white dots) from 250 neurons during the course of SD in a perfused slice. The spread of spiking and the K + wave can be seen in real-time. We recommend downloading the file and using slower playback to visualize spread of hypoxia.

**Extended Data 2 - Supporting Movie 5-1:** Supporting Movie 5-1 shows extracellular ion and O_2_ concentrations across the slice averaged over depth, as well as neuronal spiking (white dots) from 250 neurons during the course of hypoxic SD-like depolarization. The spread of spiking and the K^+^ wave can be seen in real-time. We recommend downloading the file and using slower playback to visualize the delay between initiating the simulation and the spread of the K^+^ wave.

